# Neural correlates of licking behavior modulated by target position in the striatal matrix compartment

**DOI:** 10.64898/2026.04.18.719363

**Authors:** Taishi Kimoto, Tomohiko Yoshizawa, Yuta Ishimaru, Tadashi Inui, Koichi Nakamura, Yasutaka Yawaka, Makoto Funahashi

**Author notes:** **Correspondence:** Tomohiko Yoshizawa, Oral Physiology, Department of Oral Functional Science, Faculty of Dental Medicine and Graduate School of Dental Medicine, Hokkaido University, Kita 13, Nishi 7, Kita-ku, Sapporo, Hokkaido 060-8586, Japan. These authors contributed equally (Taishi Kimoto, Tomohiko Yoshizawa).

## Abstract

The striatum is a major cortical input site of the basal ganglia and plays a critical role in the control of orofacial movements such as licking. However, how striatal activity relates to the spatial features of licking behavior remains unclear. In this study, we examined whether neural activity in the striatal matrix and striosomal compartments is associated with the spatial position of a licking target during an operant task. Head-fixed mice performed a licking task in which the target positions were varied across three spatial dimensions. Using fiber photometry in Calb1-IRES-Cre and Pdyn-IRES-Cre mice, we recorded calcium signals from matrix and striosomal neurons. Associations between neural activity, target position, and behavioral variables were quantified using linear mixed-effects modeling with cross-validation. Matrix activity prior to licking onset was primarily associated with the dorsal–ventral target position and reaction time. During licking, matrix activity was modulated by anterior–posterior and medial–lateral positions, independent of reaction time and lick count. In contrast, striosomal activity during licking was predominantly associated with the dorsal–ventral position. These findings demonstrate that neural matrix activity is systematically associated with spatial features of licking behavior, with distinct contributions before and during movement. Our results suggest that striatal matrix circuits provide task-relevant spatial signals for the control of orofacial actions.

**Significant Statement:** We show that neural activity in the striatal matrix is associated with the three-dimensional position of a licking target during an operant task. Activity prior to licking onset reflects dorsal–ventral position, whereas activity during licking is modulated by the anterior–posterior and medial–lateral positions. These findings indicate that matrix activity represents spatial aspects of licking behavior, supporting a role for the striatum in integrating motor execution with task-specific spatial information and pointing to the matrix compartment as a substrate for transforming spatial coordinates into action-specific motor commands.

## Introduction

The tongue contributes to bolus formation and transport during eating and swallowing by changing shapes and moving freely, while also enabling licking through flexible movement within and beyond the oral cavity. Licking is generated by the central pattern generators (CPGs) in the brainstem (Travers et al., 1997), but its voluntary control is modulated by the basal ganglia by sending top-down signals to the CPGs (Deniau and Chevalier, 1992; Shammah-Lagnado et al., 1992; Rossi et al., 2016; Toda et al., 2017). The striatum, the primary input site of the basal ganglia, contributes to the onset and offset of licking bouts (Bakhurin et al., 2020).

The striatum is composed of two distinct compartments: the striosome (patch) and matrix (Gerfen, 1984, 1989; Jiménez-Castellanos and Graybiel, 1989; Eblen and Graybiel, 1995; Kincaid and Wilson, 1996). These compartments differ in connectivity and function. Striosomal circuits have been implicated in cost–benefit decision-making and value-based learning (Friedman et al., 2015; Bloem et al., 2017; Yoshizawa et al., 2018), whereas the matrix is more directly linked to motor control (Graybiel and Matsushima, 2023). For example, chemogenetic inactivation of matrix neurons impairs performance in learned reach-to-grasp tasks (Lopez-Huerta et al., 2016), and recent work shows that matrix neurons exhibit early activation at locomotion onset and can promote locomotion when stimulated (Dong et al., 2025).

In our previous work, we demonstrated that matrix activity increases before the onset of lateral licking and during licking, whereas striosomal activity does not show a comparable pre-licking increase (Ishimaru et al., 2025). However, it remains unclear whether matrix activity reflects the kinematics of tongue-movement imposed by target position or instead covaries with behavioral variables such as reaction time and lick count. This question has been largely overlooked, as most studies use fixed licking targets, limiting the ability to dissociate target position related changes with licking behavior and striatal activity.

Here, we combined a head-fixed licking task with a three-dimensional manipulation of spout position to disentangle spatial coordinates and behavioral influences. Using fiber photometry, we recorded neural activity from matrix and striosomal populations in the ventrolateral striatum (VLS), a region implicated in orofacial behaviors (Bakhurin et al., 2020). We quantified the relationships between neural activity, target position, and behavior using linear mixed-effects modeling and cross-validation. Our results show that matrix activity is selectively associated with specific spout coordinates and behavioral timing, suggesting that it reflects positional information relevant to ongoing tongue movements rather than a generic change in licking output.

## Materials and Methods

### Animals

All experiments were conducted in accordance with the Hokkaido University Guidelines for Animal Care and Genetic Recombination Experiments (protocol numbers 22-0033 and 2022-057). We used seven adult male Calb1-IRES-Cre mice (129S-Calb1tm2.1(cre)Hze/J, Jackson Laboratory Cat# 028532) and six adult male Pdyn-IRES-Cre mice (129S-Pdyn(tm1.1(Cre)/Mjkr)/LowlJ, Jackson Laboratory Cat# 027958), which express Cre recombinase selectively in striatal matrix and striosomal neurons, respectively (Evans et al., 2020). Mice were housed individually under a 12 h light/dark cycle (lights on from 7:00 to 19:00), and experiments were performed during the light phase. Water intake was restricted to 1–2 ml/day for 2 days before and during the experiments. Body weight was maintained above 80% of baseline; animals falling below this threshold received supplemental water gel (1% agar).

### Behavioral task

Mice were restrained using a headplate and body tube during the behavioral task (Fig. 1A). A spout was placed near the mouth and adjusted using a micromanipulator in 0.05 mm increments. Spout-licking behavior was monitored using an infrared sensor. In each session, the spout position was varied randomly in three dimensions relative to the center of the mouth: anterior–posterior (AP), medial–lateral (ML), and dorsal–ventral (DV). Each trial began with illumination of a light-emitting diode (LED) (Fig. 1B). Upon the first lick, a 4 µl drop of 5% sucrose water was delivered. At the end of the trial, the LED was turned off, followed by a 10 ± 3 s inter-trial interval (ITI), during which the mice could lick freely. Each session lasted in 20 minutes.

**Figure 1.**
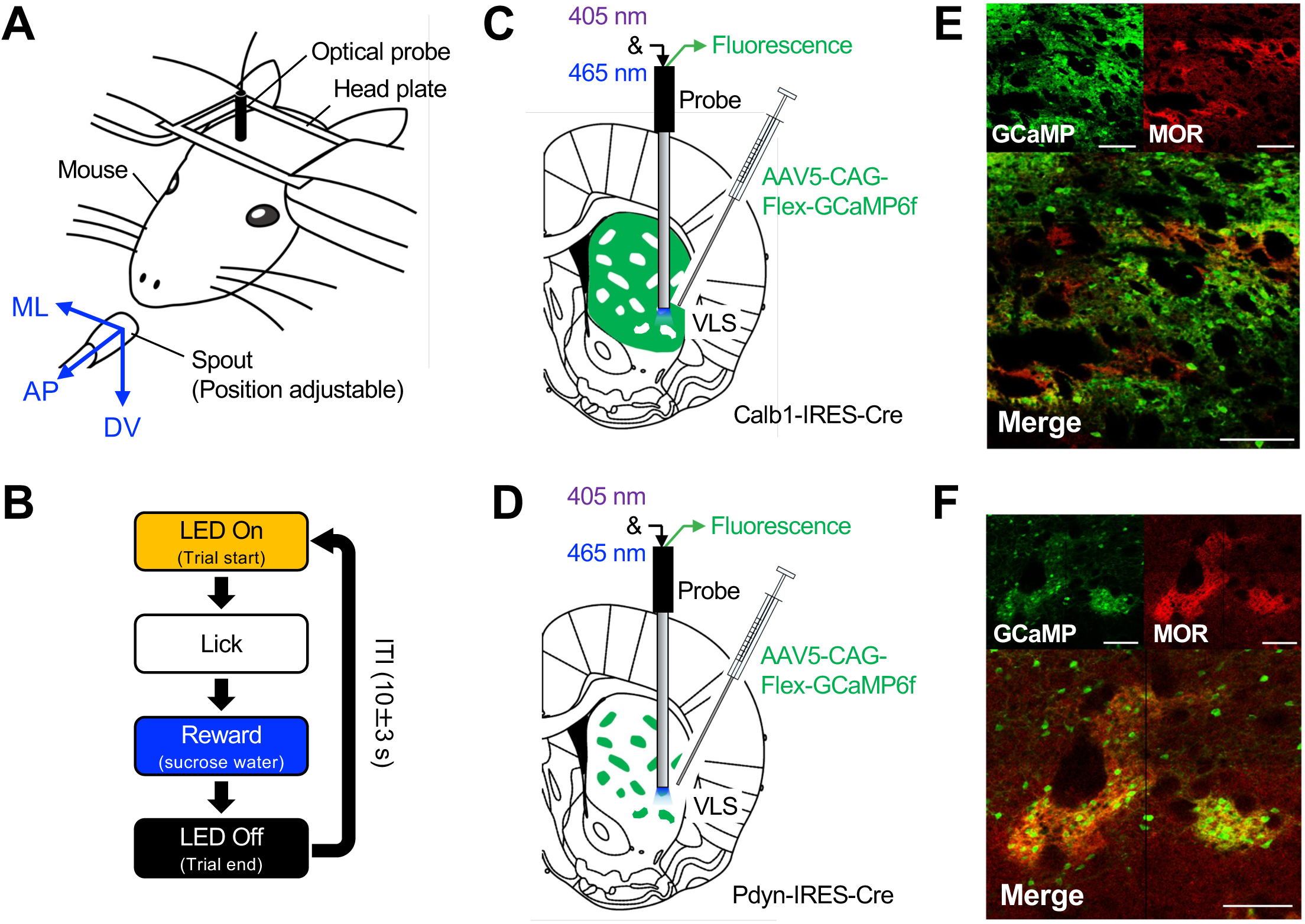
Behavioral task and fiber photometry in the striatal matrix and striosome compartments. **A.** Schematic illustration of the behavioral apparatus. The head and body of the water-deprived mouse were fixed using a metal plate and tube. A spout, adjustable in three dimensions—anterior–posterior (AP), medial–lateral (ML), and dorsal–ventral (DV)—was placed near the mouth. An optical fiber was connected to an implanted robe in the ventrolateral striatum (VLS) for fiber photometry recordings of GCaMP6f signals. **B.** Diagram of the operant conditioning task. Each trial began with illumination of a light-emitting diode (LED) in a soundproof operant box. A drop of sucrose water was delivered immediately after the mouse licked the spout. After water delivery, the LED was turned off, followed by an inter-trial interval (ITI). **C.** Schematic illustration of GCaMP fluorescence recording from matrix neurons. GCaMP6f was selectively expressed in matrix neurons via injection of AAV5.CAG.Flex.GCaMP6f into the VLS of Calb1-IRES-Cre mice. An optical probe measured calcium-dependent fluorescence of GCaMP6f excited at 465 nm. **D.** Schematic illustration of GCaMP fluorescence recording from striosomal neurons. GCaMP6f was selectively expressed in striosomal neurons via injection of AAV5.CAG.Flex.GCaMP6f into the VLS of Pdyn-IRES-Cre mice. **E.** Histological image of Cre-dependent GCaMP6f-expressing neurons in the striatum of a Calb1-IRES-Cre mouse. Scale bar: 100 µm. **F.** Histological image of Cre-dependent GCaMP6f-expressing neurons in the striatum of a Pdyn-IRES-Cre mouse. Scale bar: 100 µm

Mice were trained for 2–5 days prior to recording. During training, the spout position was gradually moved from the midline to lateral positions.

### Surgical procedures

Mice were anesthetized with isoflurane (4% induction, 1–2% maintenance) and placed in a stereotaxic frame. After exposing the skull, 400 nl of AAV.CAG.Flex.GCaMP6f (100835-AAV5, Addgene, Watertown, MA, USA) was injected into the VLS (anterior 0.26 mm, lateral 2.5 mm from bregma, and depth 3.2 mm from brain surface) using a glass capillary micropipette (Legato100, KdScientific, Holliston, MA, USA) at 40 nl/min (Fig. 1C and D).

An optical probe (diameter: 400 μm, length: 5.0 mm, R-FOC-BL400C-50NA, RWD, Guangdong, China) was implanted at a depth of 3.0 mm and secured with dental cement (Super Bond, Sun Medical, Shiga, Japan). A head plate (CF-10, Narishige, Tokyo, Japan) was affixed to the adhesive dental cement with pink dental cement (Unifast 2, GC, Tokyo, Japan). Postoperative care included analgesics (meloxicam, 1 mg/kg, s.c.) and topical antibiotics (0.1% gentamicin) as needed.

### Fiber photometry

Fiber photometry recordings were performed as previously described (Ishimaru et al., 2025; Yoshizawa and Funahashi, 2025). Excitation light at 465 nm (GCaMP) and 405 nm (isosbestic) was delivered alternately at 13.3 Hz. Emitted fluorescence was separated using dichroic mirrors (iFMC6_IE(400-410)_E1(460-490)_F1(500-540)_E2(555-570)_F2*(580-680)_S, Doric) and recorded with a photodetector.

Signals were passed through a 10 × amplifier and sampled at 1 kHz using a data acquisition system (Power1401, Cambridge Electronic Design, Cambridge, UK). The photometry signals were processed using custom-written MATLAB code (MATLAB R2018a, MathWorks, Natick, MA, USA). Signals were downsampled to 13.3 Hz for further analysis. A fitting curve was estimated and subtracted from the original signal to remove exponential and linear signal decay. The 405-nm signal was linearly fitted to the 465-nm signal, subtracted from the 465-nm channel, and divided by the 465-nm signal averaged over the session to quantify ΔF/F values. The F/F time-series trace was normalized using z-scores to account for data variability across animals and sessions. Z-scores were calculated per session using the mean and SD of each recording.

### Histology

Mice were anesthetized with pentobarbital sodium and perfused with 4% paraformaldehyde. Brains were carefully removed, post-fixed in 4% paraformaldehyde at 4◦C overnight, and transferred to a 30% sucrose/phosphate-buffered saline (PBS) solution at 4°C. Coronal sections were sliced in 30 or 100 µm thickness on a freezing microtome (REM-710; Yamato, Saitama, Japan). The 30 µm sections were processed for immunohistochemistry targeting the µ-opioid receptor (MOR, Fig. 1E and F) as previously described (Yoshizawa et al., 2018). Free-floating sections were washed in PBS for 5 min and blocked (5% normal donkey serum and 0.4% Triton X-100 in PBS) for 2 h at room temperature. Sections were incubated in rabbit anti-MOR (ab10275; Abcam) diluted 1:500 in blocking buffer for 48 h at 4°C. Sections were washed six times for 10 min in PBS and blocked for 1 h at room temperature. Sections were then incubated in donkey anti-rabbit (Alexa Fluor 594; Invitrogen) diluted 1:250 in blocking buffer for 2 h at room temperature. Sections were washed six times for 10 min in PBS and mounted on glass slides with coverslip using VECTASHIELD Mounting Medium with DAPI (Vector Laboratories, Burlingame, CA, USA). A fluorescence microscope (Eclipse Ci-L, Nikon,Tokyo, Japan) was used to analyze the tissues, and images were obtained using NIS-Elements software (NIS-Elements D, Nikon). Optical probe implantations were confirmed in all animals.

### Linear mixed-effects modeling

We fitted a linear mixed-effects model (LMM) to identify predictors of session-averaged GCaMP fluorescence. During pre-licking period (−1.0 to 0 s before the onset of first lick after LED onset), fixed effects included spout coordinates (AP, ML, and DV) and reaction time (RT, defined as the interval between LED onset and the first lick), with mouse identity included as a random effect. During licking period (0 to 1.5 s after first lick), lick count was included as an additional fixed effect. For each session (!), GCaMP fluorescence (“_!_) during pre-licking or licking periods was averaged across trials and used as a response variable. To reduce between-mouse variability, session-averaged GCaMP fluorescence and all predictor variables were mean-centered within each mouse prior to model fitting. The model was defined as:

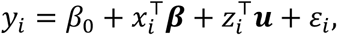

where β_0_ is the intercept, ***β*** is the regression coefficient vectors of the fixed effects, ***u*** is the vector of random effects, and ε*_i_* is the residual error term. The vectors &_!_ and (_!_ contain fixed and random effects for recording session *i*.

Model generalization was assessed using leave-one-mouse-out cross-validation. For each iteration, LMM was trained using data from all mice except one and was used to predict neural activity in the held-out mouse. Prediction performance was quantified using *R*², and statistical significance was evaluated using permutation test (10,000 permutations) in which the response variable was randomly shuffled across sessions.

### Experimental design and statistical analysis

The dataset included 5,454 behavioral and neural trials across 85 sessions in seven Calb1-IRES-Cre mice and 4,529 trials across 74 sessions in six Pdyn-IRES-Cre mice. Statistical analyses included paired or unpaired t-tests and Pearson’s correlation, as appropriate. Normality and homogeneity of variance were assessed using the Lilliefors and Bartlett tests, respectively. Statistical significance was set at *p* < 0.05.

## Results

### Effects of spout position on licking behavior

Head-fixed mice performed an operant task in which the spout position was randomly selected across behavioral sessions (Fig. 1A). For subsequent analyses, we included only trials in which no licking occurred during the 2 s preceding LED illumination. Representative licking behavior after this exclusion is shown in Fig. 2A and B. In total, 81 out of 91 trials were included. In this case, the spout was positioned at (AP, ML, DV) = (0.6, 2.0, 0.9 mm), requiring the mouse to extent its tongue out laterally to obtain the sucrose solution. Neural activity in the matrix compartment was recorded using fiber photometry in Calb1-IRES-Cre transgenic mice expressing calcium sensor GCaMP6f (Chen et al., 2013) in matrix neurons (Fig. 1C). Averaged GCaMP fluorescence across trials (Fig. 2C) showed higher fluorescence prior the onset of the first lick following LED illumination. The average fluorescence during the pre-licking period (−1.0 to 0 s before the onset of the first lick) was significantly higher than during the baseline period (−2.0 to −1.0 s) (baseline: −0.67 ± 0.024; pre-licking: −0.52 ± 0.036; *t*_80_ = −4.0, *p* = 0.00012, paired *t*-test, all fluorescence was measured using z-scores and was indicated by mean ± SEM; Fig. 2D). During the licking period (0 to 1.5 s), fluorescence was not significantly correlated with the number of licks (*r* = 0.046, *p* = 0.68; Fig. 2E).

**Figure 2.**
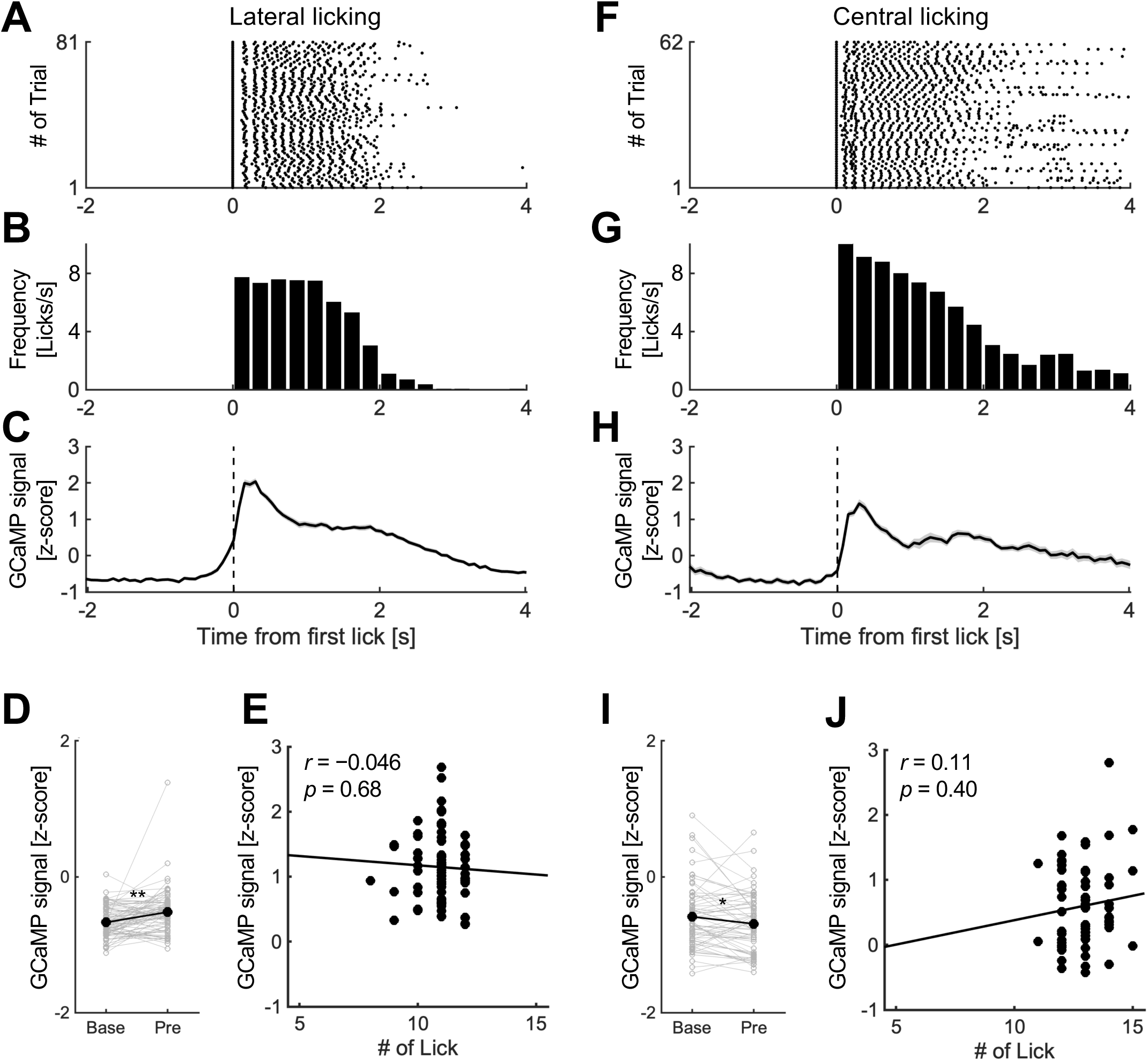
Example of neural activity in the matrix compartment during lateral and central licking. **A.** Raster plot of licking to a laterally positioned spout in a representative session. Trials are aligned to the first lick after LED onset (time 0). Black dots indicate individual lick events. **B.** Lick frequency for the session in A, computed in 250-ms bins. **C.** Session-averaged GCaMP fluorescence recorded from matrix neurons in the same session as A. Matrix neurons showed responses prior to the onset of the first lick. **D.** Mean GCaMP fluorescence during the baseline (Base; −2.0 to −1.0 s) and pre-licking (Pre; −1.0 to 0 s) periods for the session in C. Filled circles indicate trial means. Statistical analyses were performed across trials. \*\**p* < 0.01, paired *t*-test. **E.** Pearson correlation between lick count and mean GCaMP fluorescence during the licking period (0 to 1.5 s after the first lick onset) for the session in C. Each circle represents one trial; the line indicates the regression line. **F.** Same as A, but for centrally positioned spout in a representative session from the same mouse. **G.** Lick frequency for the session in F, computed in 250-ms bins. **H.** Session-averaged GCaMP fluorescence recorded from matrix neurons in the same session as F. No clear increase in activity was observed prior to lick onset. **I.** Same as D, but for the session in H. \**p* < 0.05, paired *t*-test. **J.** Same as E, but for the session in H.

In another example from the same mouse, the spout was positioned centrally at (0, 0, 1.2 mm) (Fig. 2F–H). In this case, the average fluorescence during the pre-licking period was significantly lower than baseline (baseline: −0.59 ± 0.061; pre-licking: −0.69 ± 0.056; *t*_61_ = 2.3, *p* = 0.027; Fig. 2I), and again showed no correlation with lick count (*r* = 0.11, *p* = 0.40; Fig. 2J).

To quantify the influence of spout position on licking behavior, we calculated the Pearson correlation coefficient between position and reaction time (RT). RT, defined as the interval between LED onset and the first lick after LED onset, was mean-centered within each mouse along with spatial coordinates. Using within-subject normalized values, RT showed a significant positive correlation with ML position (*r* = 0.23, *p* = 0.033) but not with AP or DV coordinates (AP: *r* = 0.15, *p* = 0.18; DV: *r* = –0.039, *p* = 0.72) (Fig. 3A-C).

**Figure 3.**
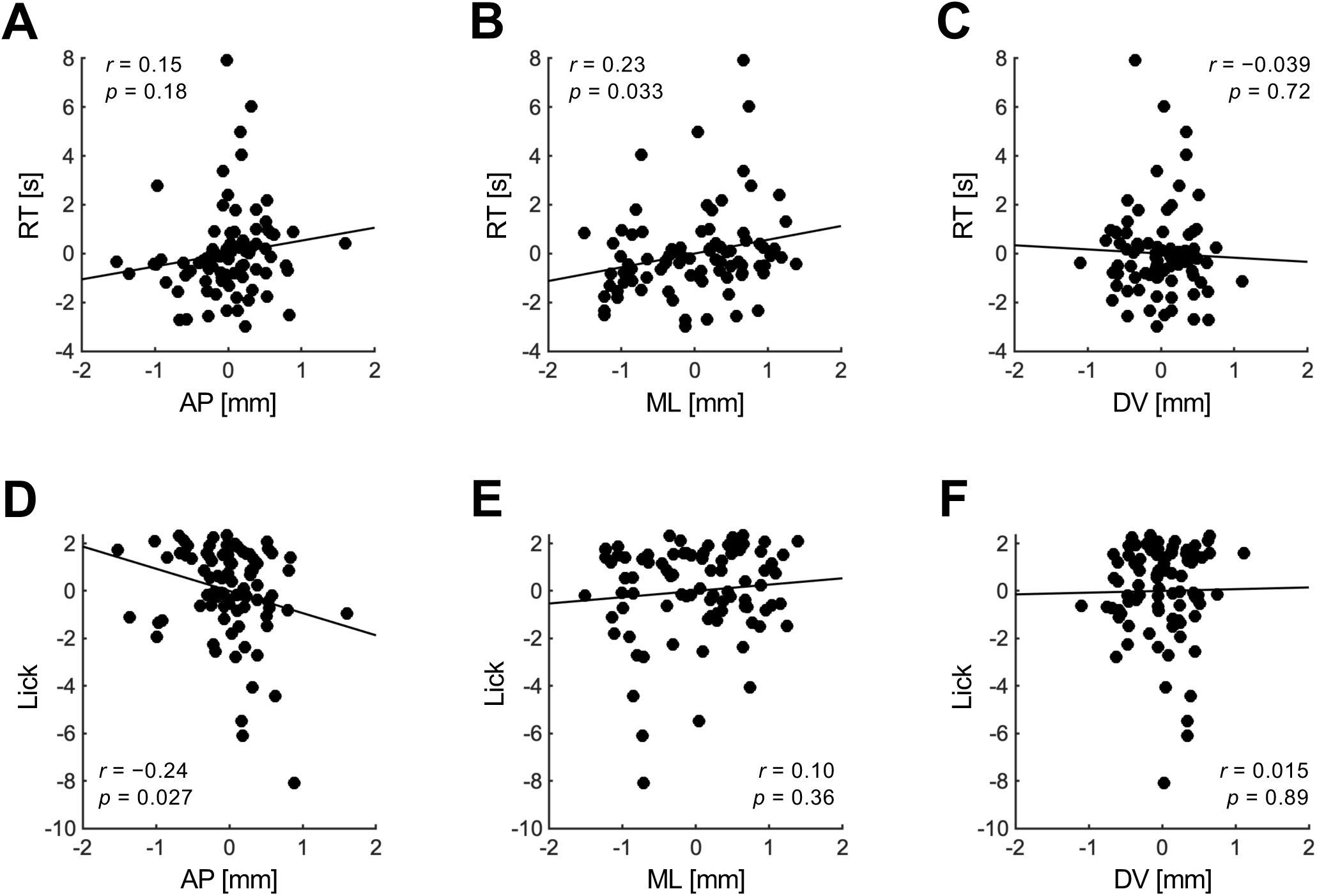
Effects of spout position on reaction time and lick number. **A.** Pearson correlation between AP coordinate and reaction time (RT). AP coordinate and RT were mean-centered within each mouse. The black line indicates the regression line. **B.** Same as A, but for ML coordinate. **C.** Same as A, but for DV coordinate. **D.** Pearson correlation between AP coordinate and lick count during the licking period. AP coordinate and lick count were mean-centered within each mouse. The black line indicates the regression line. **E.** Same as D, but for ML coordinate. **F.** Same as D, but for DV coordinate.

We next examined the correlation between spout position and the number of licks within the first 1.5 s after lick onset. After between-mouse normalization by mean-centering the data within each mouse, lick count showed a significant negative correlation with AP position (*r* = –0.24, *p* = 0.027; Fig. 3D), but not with ML or DV coordinates (ML: *r* = 0.10, *p* = 0.36; DV: *r* = 0.015, *p* = 0.89; Fig. 3E and F). These results indicate that the spout position influences response latency and licking output.

### Matrix neural activity during the pre-licking period

We first assessed the relationship between spout position and matrix neural activity during the pre-licking period using Pearson correlation. After mean-centering within each mouse, no significant correlations were observed between fluorescence and spatial coordinates (AP: *r* = 0.14, *p* = 0.21; ML: *r* = 0.17, *p* = 0.12; DV: *r* = 0.15, *p* = 0.17; Fig. 4A–C). However, session-averaged GCaMP fluorescence was positively correlated with RT (*r* = 0.42, *p* = 6.3 × 10^−05^; Fig. 4D).

**Figure 4.**
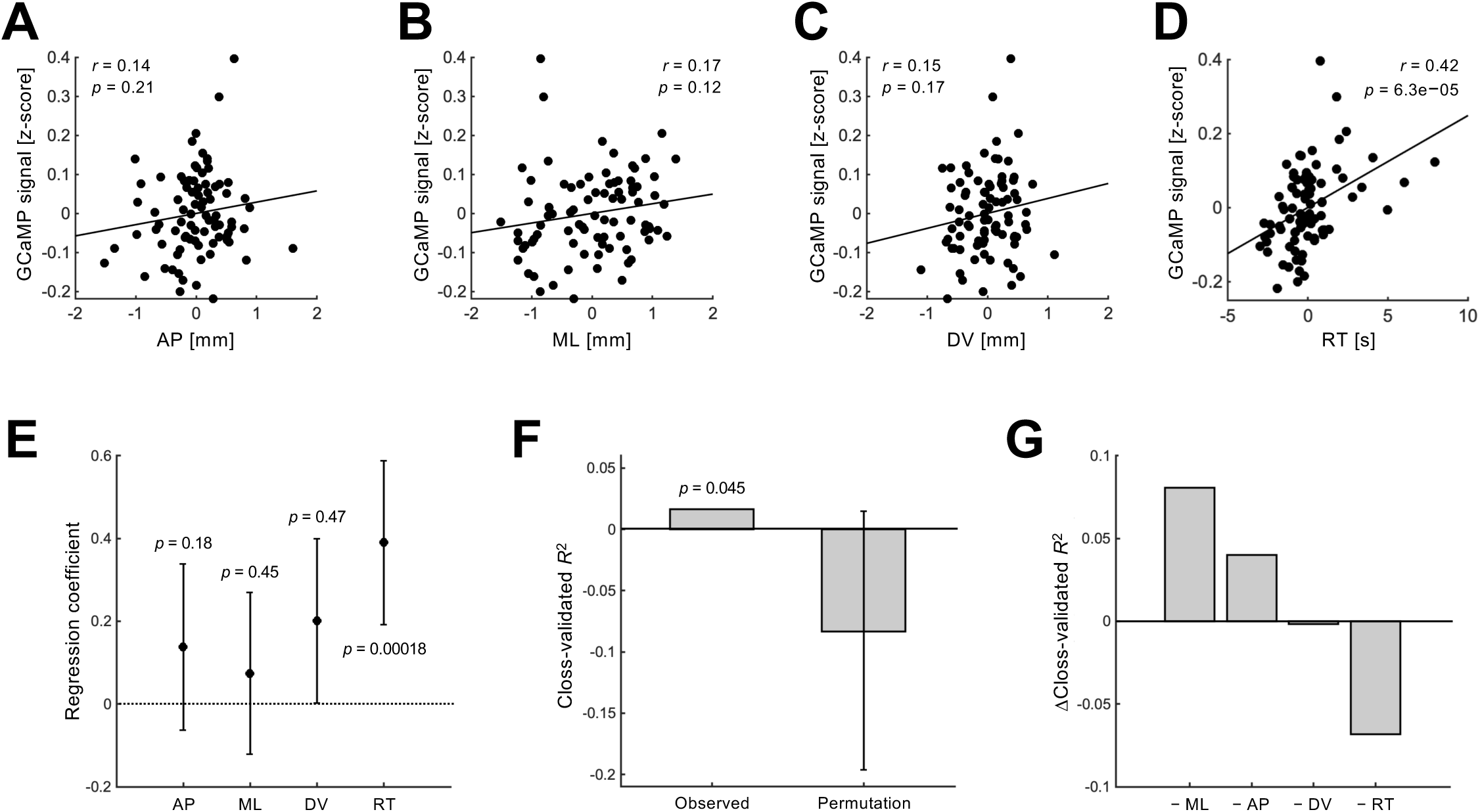
Predicting matrix neural activity during the pre-licking period from spout position and RT. **A.** Pearson correlation between AP coordinate and session-averaged GCaMP fluorescence. AP coordinate and session-averaged fluorescence were mean-centered within each mouse. The black line indicates the regression line. **B.** Same as A, but for ML coordinate. **C.** Same as A, but for DV coordinate. **D.** Pearson correlation between RT and session-averaged fluorescence. RT and session-averaged fluorescence were mean-centered within each mouse. The black line indicates the regression line. **E.** Standardized regression coefficients (*β*) from the linear mixed-effects model (LMM) predicting session-averaged GCaMP fluorescence. The model included AP, ML, DV, and RT as fixed effects and mouse identity as a random effect. Values were mean-centered within each mouse and z-scored within each session before model fitting. Filled circles and error bars indicate *β* ± 95% confidence intervals. *p* values are shown in the panel. **F.** Generalization performance of the LMM assessed using leave-one-mouse-out cross-validation. The model was trained on six mice and tested on the held-out mouse. Predictive performance (*R*^2^) was calculated on the held-out data. The observed cross-validated *R*^2^ is shown alongside a null distribution generated by permutation (10,000 iterations). Bars indicate mean cross-validated *R*^2^, and error bars for the null indicate the central 90% interval. The *p* value indicates the proportion of permuted cross-validated *R*^2^ values exceeding the observed value. **G.** Predictor-removal analysis of model generalization. One predictor was removed at a time from the full model, and the resulting change in cross-validated *R*² (Δ*R*²) was computed. Negative values indicate reduced generalization performance.

Because ML position correlated with RT (Fig. 3B) and RT showed a positive correlation with neural activity (Fig. 4D), these variables are not independent. We therefore applied a linear mixed-effects model (LMM) to predict GCaMP fluorescence. The model included AP, ML, DV, and RT as fixed effects and mouse identity as a random effect. The DV coordinate and RT were significant predictors of GCaMP fluorescence (DV: *β* = 0.20, *p* = 0.047; RT: *β* = 0.39, *p* = 0.00018; Fig. 4E), whereas AP and ML were not (AP: *β* = 0.14, *p* = 0.18; ML: *β* = 0.074, *p* = 0.45).

We next evaluated model generalization performance using leave-one-mouse-out cross-validation. Predictive performance was quantified using the coefficient of determination (*R*^2^) computed from the held-out data. The *R*^2^ was 0.016, which was significantly above chance estimated by permutation testing (*p* = 0.046; Fig. 4F). To determine the predictors that contributed to model generalization, one predictor was removed at a time from the full model and quantified the change in *R*^2^. Removing RT resulted in the largest decrease in *R*^2^ (−0.068; Fig. 4G), indicating that RT contributed most strongly to generalization. Removal DV coordinate also led to a small decrease (−0.0016), whereas removing AP or ML increased *R*^2^ (AP: +0.040; ML: +0.081), suggesting that these variables did not provide additional information for generalization.

### Matrix neural activity during the licking period

We next analyzed matrix neural activity during the licking period (0–1.5 s after the onset of the first lick). Pearson correlation analysis revealed significant positive correlations between session-averaged GCaMP fluorescence and AP (*r* = 0.44, *p* = 2.1× 10^−05^) and ML (*r* = 0.38, *p* = 3.7 × 10^−04^), and a negative correlation with DV (*r* = −0.25, *p* = 0.022; Fig. 5A–C). The session-averaged GCaMP fluorescence was also significantly correlated with RT (*r* = 0.37, *p* = 5.0 × 10^−04^) and lick count (*r* = −0.27, *p* = 0.014; Fig. 5D and E).

**Figure 5.**
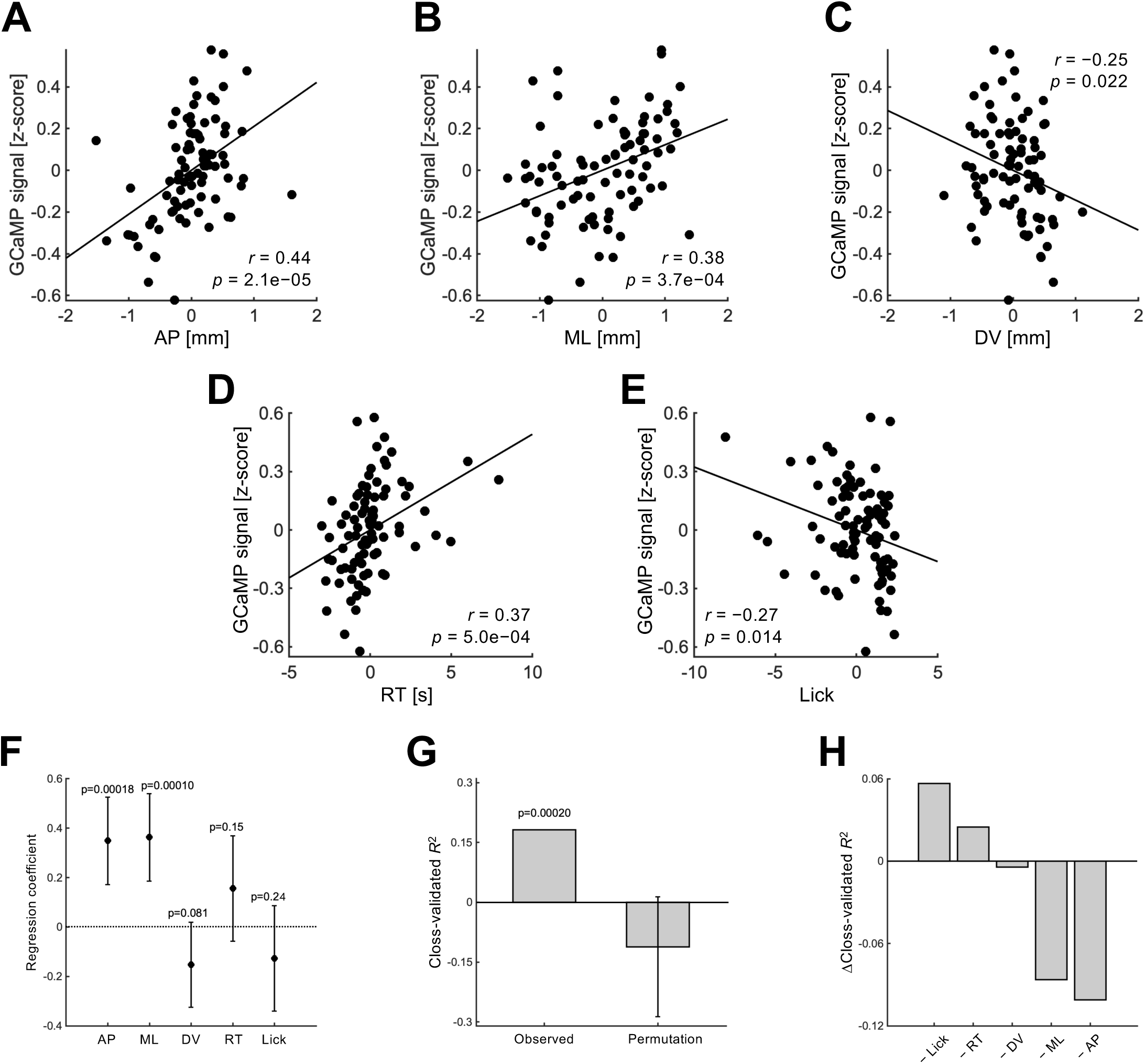
Predicting matrix neural activity during the licking period from spout position, RT and lick number. **A.** Pearson correlation between AP coordinate of the spout and session-averaged GCaMP fluorescence. AP coordinate and session-averaged fluorescence were mean-centered within each mouse. The black line indicates the regression line. **B.** Same as A, but for ML coordinate. **C.** Same as A, but for DV coordinate. **D.** Pearson correlation between RT and session-averaged fluorescence. RT and session-averaged fluorescence were mean-centered within each mouse. The black line indicates the regression line. **E.** Same as E, but for lick number. **F.** Standardized regression coefficients (*β*) from the LMM predicting session-averaged GCaMP fluorescence. The model included the AP, ML, DV, RT, and lick number as fixed effects, and mouse identity as a random effect. Values were mean-centered within each mouse and z-scored within each session before model fitting. Filled circles and error bars indicate *β* ± 95% confidence intervals. *p* values are shown in the panel. **G.** Generalization performance of the LMM for neural activity assessed using leave-one-mouse-out cross-validation. The model was trained on data from six mice and tested on the held-out mouse. Predictive performance (*R*^2^) was calculated on the held-out data. The observed cross-validated *R*^2^ is shown alongside a null distribution generated by permutation (10,000 iterations). Bars indicate mean cross-validated *R*^2^, and error bars for the null indicate the central 90% interval. The *p* value indicates the proportion of permuted cross-validated *R*^2^ values exceeding the observed value. **H.** Predictor-removal analysis of model generalization. One predictor was removed at a time from the full model, and the resulting change in cross-validated *R*² (Δ*R*²) was computed. Negative values indicate reduced generalization performance.

As spout coordinates, RT, and lick count were not independent of each other (Fig. 3), we applied an LMM to evaluate their independent contributions to neural activity during licking. The model included AP, ML, DV, RT, and lick count as fixed effects, and mouse identity as a random effect. Only AP and ML were significant predictors of GCaMP fluorescence (AP: *β* = 0.35, *p* = 1.8 × 10^−04^; ML: *β* = 0.36, *p* = 1.0 × 10^−04^; Fig. 5F). In contrast, DV, RT, and lick count were not significant (DV: *β* = −0.15, *p* = 0.081; RT: *β* = 0.16, *p* = 0.15; lick: *β* = −0.13, *p* = 0.24).

Cross-validation yielded an *R*² of 0.18, significantly above chance (*p* = 2.0 × 10^−04^; Fig. 5G). Removing AP, ML, or DV reduced model performance (AP: −0.10; ML: −0.086; DV: −0.042; Fig. 5H), whereas removing RT or lick count increased *R*² (RT: +0.025; lick: +0.057). These results indicate that spatial variables, particularly AP and ML, contributes to model generalization during licking.

### Striosomal neural activity during the licking period

We recorded neural activity in the striosomal compartment during the operant task using fiber photometry in Pdyn-IRES-Cre transgenic mice expressing GCaMP6f in striosomal neurons in a Cre-dependent manner (Fig. 1D). Fig. 6A–C show lateral licking behavior and striosomal neural activity recorded from a representative mouse. The mouse extended its tongue laterally to drink sucrose solution from the spout with coordinates of 1.0 mm anteriorly, 3.05 mm laterally, and 0.6 mm ventrally from the center of the oral fissure. Fig. 6D and E show an example of central licking, and Fig. 6F shows GCaMP fluorescence from the same session. These data were recorded from the same mouse, as shown in Fig. 6A–C. During central licking, the spouts were positioned at (AP, ML, DV) = (0.2, 0, 0.8) mm. In both cases, no clear increase in neural activity was observed prior to lick onset, consistent with previous findings (Ishimaru et al., 2025).

**Figure 6.**
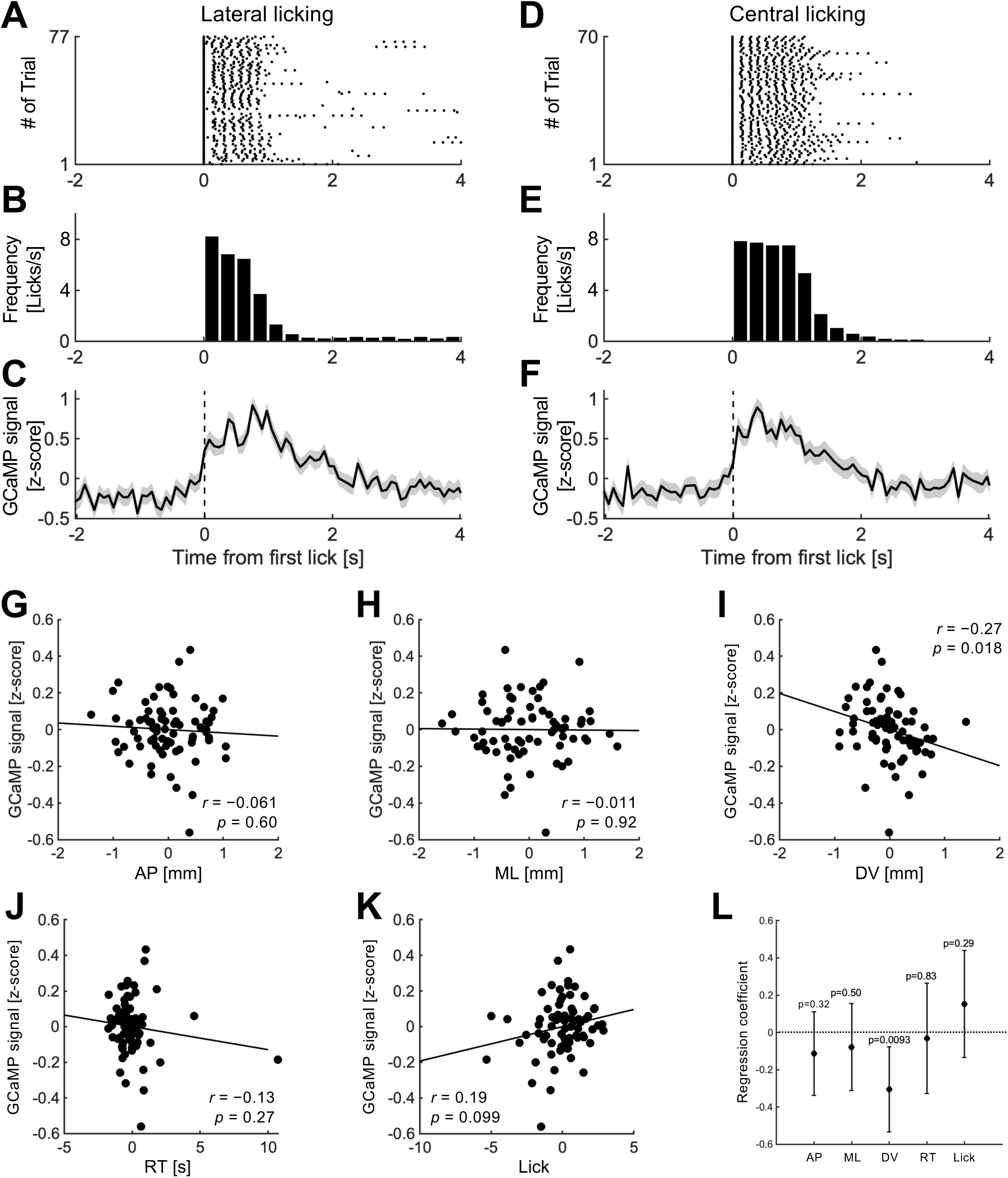
Neural activity in the striosome compartment. **A.** Raster plot of licking to a laterally positioned spout in a representative session. Trials are aligned to the first lick after LED onset (time 0). Black dots indicate individual lick events. **B.** Lick frequency for the session in A (250-ms bins). **C.** Session-averaged GCaMP fluorescence recorded from striosomal neurons in the same session as A. **D.** Same as A, but for a centrally positioned spout in a representative session from the same mouse. **E.** Lick frequency for the session in D (250-ms bins). **F.** Session-averaged GCaMP fluorescence recorded from striosomal neurons in the same session as D. **G.** Pearson correlation between AP coordinate and session-averaged GCaMP fluorescence during the licking period. AP coordinate and session-averaged fluorescence were mean-centered within each mouse. The black line indicates the regression line. **H.** Same as G, but for ML coordinate. **I.** Same as G, but for DV coordinate. **J.** Pearson correlation between RT and session-averaged fluorescence. RT and session-averaged fluorescence were mean-centered within each mouse. The black line indicates the regression line. **K.** Same as J, but for lick count. **L.** Standardized regression coefficients (*β*) from the LMM predicting session-averaged GCaMP fluorescence during the licking period. The model included AP, ML, and DV, RT, and lick count during the licking period as fixed effects, and mouse identity as a random effect. Values were mean-centered within each mouse and z-scored within each session before model fitting. Filled circles and error bars indicate *β* ± 95% confidence intervals. *p* values are shown in the panel.

We therefore focused on the licking period. After mean-centering within each mouse, session-averaged GCaMP fluorescence was not significantly correlated with AP or ML coordinates (AP: *r* = −0.061, *p* = 0.60; ML: *r* = −0.011, *p* = 0.92; Fig. 6G and H), but showed a significant negative correlation with DV (*r* = −0.27, *p* = 0.018; Fig. 6I). No significant correlations were found with RT or lick count (RT: *r* = −0.13, *p* = 0.27; Lick: *r* = 0.19, *p* = 0.099; Fig. 6J and K).

LMM analysis revealed that only DV was a significant predictor of GCaMP fluorescence (*β* = −0.31, *p* = 0.0093; Fig. 6F), whereas all other variables were not (AP: *β* = −0.11, *p* = 0.32; ML: *β* = −0.079, *p* = 0.50; RT: *β* = −0.031, *p* = 0.83; Lick: *β* = 0.15, *p* = 0.29). Cross-validation yielded an *R*² of −0.13, not different from chance (*p* = 0.60), indicating poor generalization across mice.

## Discussion

In this study, we investigated how striatal neural activity reflects licking targets. The main findings are: (1) spout position influenced both RT and the number of licks during licking period; (2) matrix neural activity during pre-licking period correlated with RT and DV coordinate; and (3) matrix neural activity during licking was associated with AP and ML coordinates, whereas striosomal neural activity did not show such association.

### Effects of spout position on licking behavior in mice

Licking behavior is essential for drinking and eating. It is one of the most widely used behavioral measures in neuroscience, physiology, and psychology. Previous studies have focused on licking frequency in response to a conditioned stimulus in classical conditioning paradigms or lateralized licking to investigate stimulus discrimination or decision-making processes (Oyama et al., 2010; Cohen et al., 2012; Yoshizawa et al., 2018; Handa et al., 2021; Lee and Sabatini, 2021; Funamizu et al., 2023; Yoshizawa and Funahashi, 2025). However, these studies have not systematically examined how spout position affects licking behavior.

By varying spout position in three dimensions, we found that the distance between oral fissure and spout along the ML axis was positively correlated with RT, whereas the AP axis was negatively correlated with the number of licks during the licking period. These effects reflect biomechanical constraints: more distant targets require greater tongue extension, increasing RT and reducing lick count. These results imply that the distance between oral fissure and spout must be held constant within each behavioral session to ensure consistent performance.

### Pre-lick neural signals and their behavioral role

Consistent with our previous work, we observed neural responses prior to lateral licking onset in the matrix compartment (Ishimaru et al., 2025). As previous study showed that these responses are stronger for licks at spout located ipsilateral to the recording hemisphere, we recorded from the right hemisphere and trained the mice to lick a spout positioned on the right side of the oral fissure. While simple Pearson correlations did not detect significant correlations between matrix neural activity and spout coordinates during pre-licking period, LMM identified a significant effect of the DV coordinate on neural activity. This discrepancy indicates that the effect of DV coordinate can be captured only when multiple behavioral variables are considered simultaneously.

In addition, we also observed a significant positive association between neural activity and RT. Body movements are often considered as reflexive movements, which are triggered by external sensory stimuli and voluntary movements, and initiated under intentional control. Reflexive movements typically exhibit very short reaction times as only few synaptic relays are involved in the sensorimotor pathways. On the other hand, voluntary movements require longer reaction times because of the additional processing required for motor planning and decision-making (Kurtzer, 2014). The strong association between pre-lick neural activity in the matrix compartment with RT suggests that pre-lick neural activity reflects preparatory processes linked to voluntary control rather than fast reflexive responses. It further indicates that neurons in the matrix compartment contribute to the preparation and execution of orofacial actions by encoding preparatory demand levels before movement onset. This role is consistent with previous works indicating that matrix neurons participate in the selection and execution of flexible goal-directed behaviors (Barto, 1995; Doya, 2002).

### Interpretation of neural correlates of licking target position

Our LMM indicated that matrix neural activity during licking was positively associated with AP and ML coordinates, whereas no such associations were observed in the striosome compartment. These results suggest that matrix neural activity is more strongly related to licking movement direction than striosomal neural activity. Given the presence of head-direction cells in the striatum (Wiener, 1993; Mizumori et al., 2000), the activity of individual neurons in the matrix compartment may be tuned to the direction of licking movements. Further studies are required to determine whether individual matrix neurons are selectively tuned to a specific licking direction or respond to multiple directions.

The matrix compartment contains approximately equal proportion of direct spiny projection neurons (dSPNs) expressing dopamine D1 receptors or indirect spiny projection neurons (iSPNs) expressing dopamine D2 receptors, whereas the striosome is enriched in dSPNs (Fujiyama et al., 2011; Miyamoto et al., 2018). A previous study reported that optogenetic activation of iSPNs, without accounting for striatal compartments, suppresses contralateral licking and promotes ipsilateral licking (Lee and Sabatini, 2021). Our results show that matrix neural activity was higher during ipsilateral compared to central licking. These observations suggest that increased matrix activity during ipsilateral licking may reflect, at least in part, the engagement of iSPN-related circuits.

Consistent with a potential role of iSPN-related circuits, dopamine deficiency in the striatum−a hallmark of Parkinson’s disease (PD)−has been shown to impair licking movements in rodents (Skitek et al., 1999; Ciucci et al., 2011). Moreover, hemi-PD mouse models display lateral deviations of tongue protrusions toward the affected side (Chen et al., 2019). Because striatal dopamine depletion suppresses dSPN activity while enhancing iSPNs activity, iSPN-related circuits are expected to be enhanced under parkinsonian conditions (Gerfen, 2022). In contrast, D2 receptor antagonists, such as chlorpromazine, induce parkinsonian motor symptoms by blocking nigrostriatal D2 signaling (Sykes et al., 2017). Such pathway imbalance may contribute to the lateralized tongue movements observed in PD. Further research is needed to determine how the relative activity of dSPNs and iSPNs in the matrix compartment is altered under parkinsonian conditions, and how this imbalance contributes to impaired orofacial motor control.

## Author Contributions

T.Y. designed the study; T.Y., T.K. and Y.I. performed the study; T.Y. and T.K. analyzed the data; T.Y., T.I, K.N., Y.Y. and M.F. supervised the study; T.Y. wrote the paper; T.Y. acquired the funding.

## Acknowledgements

This work was supported by JSPS KAKENHI Grant Numbers JP22K15247 and JP25K09843, the Northtec Foundation, and a generous research support from Hokkaido University for Young Researchers. We would like to thank Editage (www.editage.jp) for English language editing.

